# Kinetics-based Inference of Environment-Dependent Microbial Interactions and Their Dynamic Variation

**DOI:** 10.1101/2022.08.27.505268

**Authors:** Hyun-Seob Song, Na-Rae Lee, Aimee K. Kessell, Hugh C. McCullough, Seo-Young Park, Kang Zhou, Dong-Yup Lee

## Abstract

Microbial communities in nature are dynamically evolving as member species change their interactions subject to environmental variations. Accounting for such context-dependent dynamic variations in interspecies interactions is critical for predictive ecological modeling. In the absence of generalizable theoretical foundations, we lack a fundamental understanding of how microbial interactions are driven by environmental factors, significantly limiting our capability to predict and engineer community dynamics and function. To address this issue, we propose a novel theoretical framework that allows us to represent interspecies interactions as an explicit function of environmental variables (such as substrate concentrations) by combining growth kinetics and a generalized Lotka-Volterra model. A synergistic integration of these two complementary models leads to the prediction of alterations in interspecies interactions as the outcome of dynamic balances between positive and negative influences of microbial species in mixed relationships. This unique capability of our approach was experimentally demonstrated using a synthetic consortium of two *Escherichia coli* mutants that are metabolically dependent (due to an inability to synthesize essential amino acids), but competitively growing on a shared substrate. The analysis of the *E. coli* binary consortium using our model not only showed how interactions between the two amino acid auxotrophic mutants are controlled by the dynamic shifts in limiting substrates, but also enabled quantifying previously uncharacterizable complex aspects of microbial interactions such as asymmetry in interactions. Our approach can be extended to other ecological systems to model their environment-dependent interspecies interactions from growth kinetics.

**IMPORTANCE:** Modeling of environment-controlled interspecies interactions through separate identification of positive and negative influences of microbes in mixed relationships is a new capability that can significantly improve our ability to understand, predict, and engineer complex dynamics of microbial communities. Moreover, robust prediction of microbial interactions as a function of environmental variables can serve as valuable benchmark data to validate modeling and network inference tools in microbial ecology, the development of which has often been impeded due to the lack of ground truth information on interactions. While demonstrated against microbial data, the theory developed in this work is readily applicable to general community ecology to predict interactions among microorganisms such as plants and animals, as well as microorganisms.

## INTRODUCTION

Microbial communities play pivotal roles in maintaining human and animal health, plant productivity, and ecosystem services (1-4). Increasing efforts are being dedicated towards maximizing their beneficial roles in natural systems or creating new industrial applications (5). However, control and design of microbial community dynamics and function is a challenging task, primarily due to higher-order or emergent properties that are not observable from individual species in isolation but arise through nonlinear interspecies interactions (6, 7). Therefore, rational design of microbial communities or consortia requires a fundamental knowledge of microbial interactions as a mechanistic linkage between the environment and the community compositions and function, necessitating the employment of predictive mathematical models as indispensable tools (8-14).

Development of accurate models of microbial communities that are commonly subject to environmental variations is truly complicated by the following intrinsic ecological aspects. First, microorganisms in a community build *dynamic* interactions that cannot effectively be represented by a rigid network with fixed structure (15, 16). Rather, microbial communities keep reorganizing interaction networks in response to biotic or abiotic perturbations or through adaptation to long-lasting environmental changes. Second, microorganisms often build *mixed* relationships by exerting *both* promotive and inhibitive impacts on the growth of their partners/neighbors (17, 18). Individual identification of these simultaneously acting positive and negative interactions is critical because community dynamics is mainly driven by the balances between all counteracting impacts among member species (19). The lack of capability to account for these key properties of microbial interactions limits our ability to predict and engineer microbial community dynamics and functions.

Despite rapid progress in microbiome science, we still do not know how to identify environment-controlled dynamic variation in interspecies interactions addressed above. Three major branches of microbial interaction modeling include (20, 21): (i) network inference, (ii) metabolic network modeling, and (iii) kinetic modeling. Network inference is widely used for modeling microbial interactions to identify interaction networks based on correlative relationships among microbial populations (22-25), parameter identification through regression (26-28), or a prescribed set of rules or hypotheses (21). The resulting networks represent interspecies interactions as single constant metrics, therefore being unable to describe dynamic variations in interactions nor identify the balances among counteracting individual impacts in mixed relationships. As an exception, the approach termed MIIA (Minimal Interspecies Interaction Adjustment) (15, 16) uniquely enables predicting context-dependent interactions due to the changes in memberships, which however has not been extended to address the environmental impacts. In contrast with such data-driven network inference methods, metabolic network and kinetic modeling can account for both positive and negative interactions based on cross-feeding of small molecules (essential for growth) or competition for shared substrates/nutrients among species; in theory, kinetic models can additionally simulate their dynamic variations. While more mechanistic than network inference, these methods cannot quantify the magnitude or even the sign of net interactions.

In this work, we fill these gaps by proposing a novel theoretical framework that enables a quantifiable, mechanistic representation of the dynamic linkage between microbial interactions and the environment. For this purpose, we synergistically integrate two complementary modeling frameworks to overcome their own limitations: a generalized Lotka-Volterra (gLV) model (29) and population growth kinetics. Like other network inference approaches, a typical gLV model with a focus on pairwise interactions is constructed based on an implicit assumption of constant interactions. We relax this assumption by representing interaction coefficients in the gLV model as a function of environmental variables (i.e., concentrations of cross-fed metabolites and shared substrates) described in microbial growth kinetics, which is termed here kinetics-based inference of dynamic variation in microbial interactions (KIDI). The resulting functional representation of interactions by KIDI enables not only quantifying their dynamic variation as environmental conditions change, but also individually identifying negative and positive influences among species in mixed relationships. The prediction of KIDI was demonstrated accurate through a coordinated design of experiments using a binary consortium composed of tyrosine and tryptophan auxotrophic mutants of *Escherichia coli* (30) so that they both compete and/or cooperate depending on environmental conditions.

## RESULTS

### Formulation of a conceptual model for understanding environment-dependent interactions

For illustration of the concept of KIDI, we consider a hypothetical consortium composed of two members where species 1 (X_1_) and species 2 (X_2_) cooperate by cross-feeding 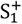 and 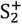 each other but compete for the shared metabolite S^−^ (the center circle in **Fig. 1**). Growth kinetics for the *i*^*th*^ species (X_i_) (that requires two substrates 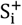 and S^−^ for growth) can be represented, e.g., using a double Michaelis-Menten equation as follows:

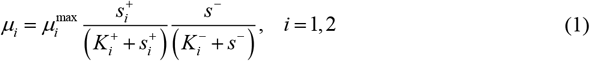

where μ_*i*_ [1/h] is the specific growth rate of X_i_, 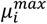 is the maximal specific growth rate, 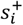 and *s*^−^ [g/l] are the concentrations of 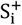 and S^−^, and 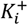 and 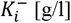 are half-saturation constants associated with the consumption of 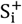 and S^−^, respectively. As inferable from growth kinetics in Eq. (1), the mixed relationship (i.e., competition and cooperation) between X_1_ and X_2_ when both substrates are limiting can turn into diverse forms of interactions as environmental conditions change. When S^−^ is present in excess (therefore, no competition is necessary) but 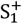 and 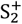 are limiting, for example, their relationship is predominantly cooperative (where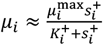). As the opposite case, if both 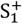 and 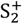 are excessive in the environment (so no partners are needed to acquire them) while S^−^ is limiting, their relationship is governed by competition (where 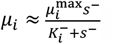). Likewise, one can assume many other different scenarios where their relationships turn into competition, cooperation, amensalism, commensalism and even neutrality, as illustrated in **Fig. 1**.

**Fig. 1.**
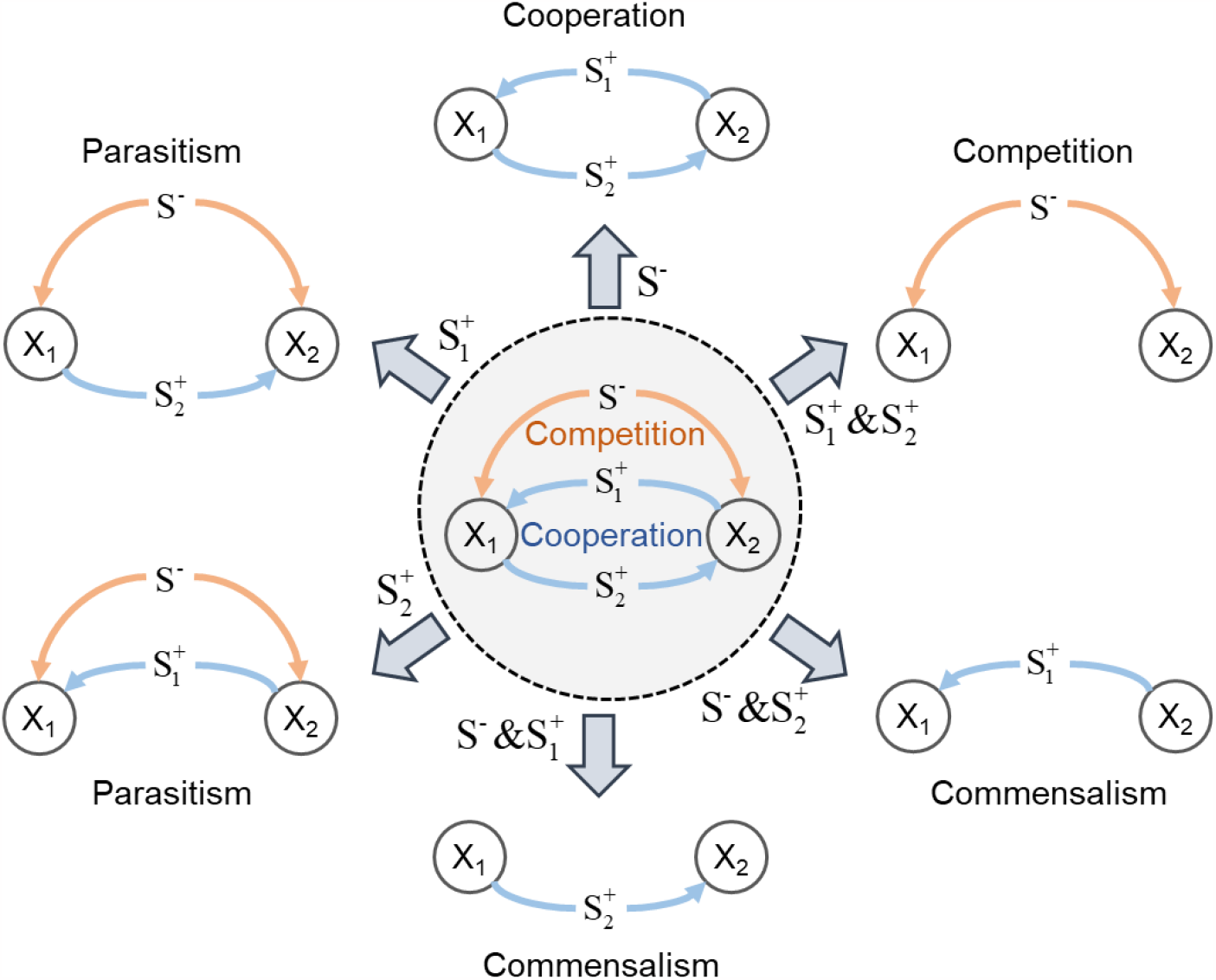
Conceptual illustration of context-dependent microbial interactions in a binary consortium dictated by the environmental contexts. Two species X_1_ and X_2_ compete for the substrate S^−^ but cooperate by cross-feeding metabolite 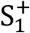 and 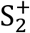 (center panel). This mixed relationship between X_1_ and X_2_ diverge into six different types of interactions by excessive addition of specific substrates 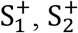, and/or S^−^. Symbols next to the arrows denote the substrate(s) excessively added to the environment. (**c**) Model microbial consortium using tyrosine or tryptophan auxotrophic *E. coli* strains. Glucose is sole carbon source for both *E. coli* strains. GLC, Tyr and Trp denote glucose, tyrosine, and tryptophan, respectively.

### Representation of interaction parameters as a function of environmental variables

To model such environment-dependent microbial relationships, we derived a general form of interaction coefficients as a function of environmental variables by integrating growth kinetics and a gLV model. As described in detail in **Materials and Methods**, our formula (KIDI) represents interaction coefficients of species in the mixed relationship as a sum of positive and negative parts, i.e.,

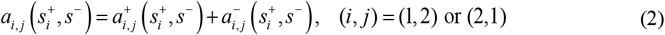

where 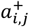 and 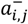 denote the positive and negative influence of X_j_ on the growth rate of X_i_, which are defined as follows:

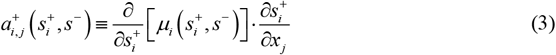

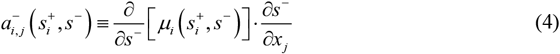

The positive influence of X_j_ on the growth rate of 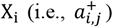 is represented by the two subsequent terms on the right-hand side of Eq. (3): (i) the impact of the change in the population size of X_j_ on the concentration of the cross-fed substrate 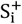 (as denoted by 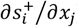 and (2) the subsequent impact of the change in 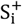 on the growth rate of the *i*^*th*^ species (i.e., *μ*_*i*_) (as denoted by 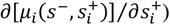. The negative impact of X_j_ on the growth rate of 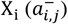 in Eq. (4) can be interpreted in a similar fashion.

The derivative terms on the right-hand side of Eqs. (3) and (4) are fully identifiable from reaction stoichiometry and kinetics. In the case of using a double Monod kinetics, for example, incorporation of Eq. (1) into Eqs. (3) and (4) yields 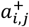 and 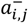 as follows:

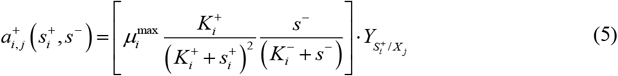

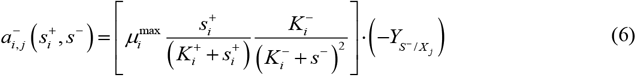

where 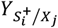and 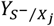 denote the stoichiometric relationships between the changes in substrate and biomass concentrations associated with X_j_, i.e., 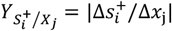 and 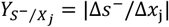 (see **Materials and Methods**).

To identify net interactions between two species with mixed relationships, we further defined a normalized interaction parameter *γ*_*i,j*_ as follows:

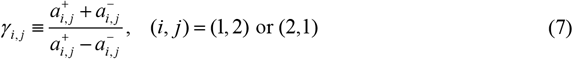

Once 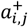 and 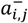are identified from Eqs. (3) and (4), the parameter *γ*_*i,j*_ is readily calculatable by Eq. (7). The parameter *γ*_*i,j*_ ranges from -1 to 1 to represent positive influences of species *j* on species *i* when greater than 0 and negative impacts when less than 0, respectively, consequently allowing us to conveniently quantify the relative dominance of inhibition vs. promotion in mixed interactions. The parameter *γ*_*i,j*_ complements *a*_*i,j*_, rather than replaces it, in that the magnitude of interactions cannot be determined by *γ*_*i,j*_, but by the original interaction parameter, *a*_*i,j*_. In this regard, *γ*_*i,j*_ provides an additional complementary explanation of dynamic changes in interspecies interactions. Therefore, all these parameters, including *a*_*i,j*_ defined in Eqs. (2) to (4) and *γ*_*i,j*_, sufficiently characterize the dynamic variation of interactions between X_1_ and X_2_ based on co-culture growth data as demonstrated in the following sections.

### Identification of kinetics and stoichiometry via data fit

For experimental demonstration of the mathematical formulation derived in the previous section, we constructed a synthetic consortium composed of two *E. coli* auxotrophic mutants that can cooperatively cross-feed small molecules, while competitively growing on a common substrate (31). Among 14 amino acid auxotrophic mutants, we chose tryptophan and tyrosine auxotrophic mutants by considering the bioenergetic cost for the synthesis of amino acids based on a previous study in the literature (30). This consortium is considered an ideal, simplest model system for studying environment-dependent dynamic variations in microbial interactions. Due to its exact correspondence to the hypothetical consortium in **Fig. 1**, we denote two *E. coli* mutant strains ΔtrpC and ΔtyrA by X_1_ and X_2_; glucose, tryptophan, and tyrosine by 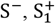, and 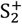, respectively.

Using these two strains, we performed growth experiments under diverse culture conditions: two individual batch experiments using X_1_ (**Fig. 2A**) and X_2_ (**Fig. 2B**), respectively, and two sets of co-culture experiments (**Figs. 2C** and **2D**). The top panels in **Figs. 2C** and **2D** denote co-growth experiments in batch cultures, while the middle and bottom panels denote semi-batch modes, where we added glucose feedbeads (FBs) at time 7.5 and 10 hours, respectively, to induce dramatic changes in interspecies interactions during co-growth. Other differences in co-culture conditions in **Figs. 2C** and **2D** include initial concentrations of 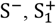, and 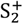, and the number of added FBs (see **Supp Datasheet**). We measured optical density (OD) at an absorbance of 600 nm as a metric of cell density and determined the OD for each strain in the consortium through additional analysis using quantitative PCR (qPCR). The OD profiles in **Fig. 2D** denote the combined population change of both strains, i.e., X_1_ + X_2_.

**Fig. 2.**
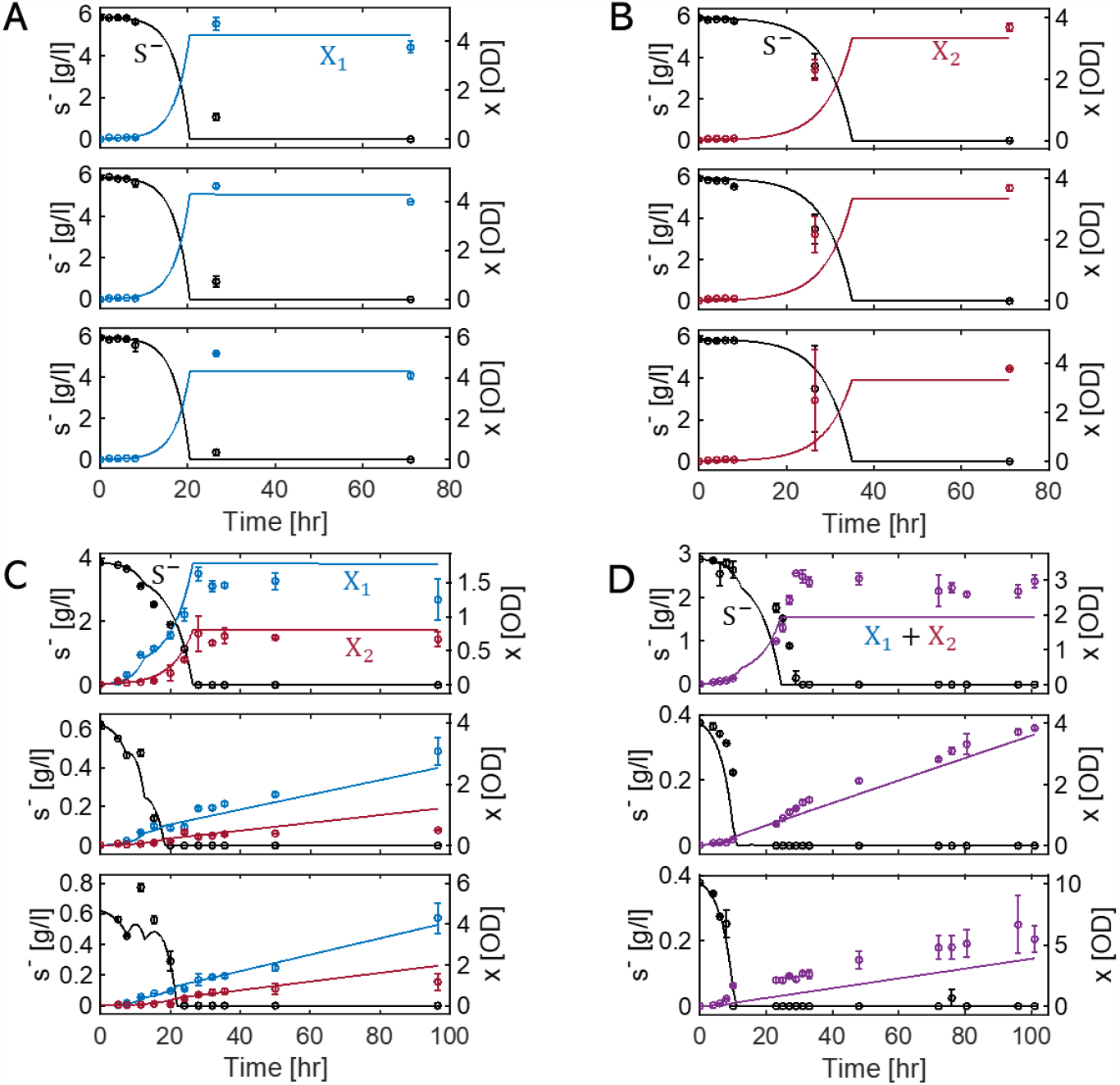
Experimental data and model simulations for the growth of two *E. coli* mutant strains in axenic and binary culture conditions: **A** and **B** are cultures of tryptophan auxotrophic and tyrosine auxotrophic *E. coli* mutants (X_1_ and X_2_), respectively; **C** and **D** are co-cultures with two auxotrophs in batch and semi-batch cultures. Detailed culture conditions for the twelve panels are provided in **Table S2**. Circles and lines denote the experimentally measured values and simulation results, respectively. Black line denotes simulation results for glucose concentration (S^−^), and the lines in blue, red, and purple are simulated population densities of X_1_, X_2_, and X_1_ + X_2_. **Figs. A, B**, and **C** show data fitting to determine model parameters, while the results in **Fig. D** validate model predictions.

Based on the four datasets in **Fig. 2**, we constructed a dynamic co-growth model of X_1_ and X_2_ to determine associated kinetics and stoichiometry, key information required for quantifying interspecies interactions parameters (*a*_*i,j*_ and *γ*_*i,j*_) in Eqs. (2) to (7). The dynamic co-growth model is composed of five mass balance equations for X_1_, X_2_, S^−^, 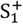, and 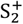 . We determined stochiometric and kinetic parameters using three subsets of data in **Figs. 2A – 2C** and validated the model against the remaining one (in **Fig. 2D**) that was not used for model identification. The robust consistency between simulated and measured data in **Fig. 2D**, as well as those in **Figs. 2A – 2C**, indicates the acceptability of using the identified model parameters in inferring interaction coefficients. We provide the full list of model equations with parameter values in **Table 1**; the culture conditions in **Table S1**.

**Table 1.** Model equations with kinetic parameters and stoichiometric coefficients determined through the model fit to experimental data collected under various limiting conditions. R_i_ is the stoichiometric growth reaction for X_i_, and 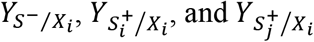denote the stoichiometric coefficients for S^−^, 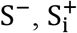, and 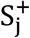 associated with the growth of X_i_. *s*^−^, 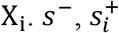, and 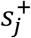 respectively denote concentrations of 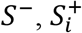, and 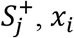, *x*_*i*_ is the population density of X_i_, μ_*i*_ is the specific growth rate of X_i_, *k*_*d,i*_ is the death rate of X_i_, and *q*_*s*−_ is the substrate releasing rate from FBs in glucose-limited semi-batch cultures (i.e., 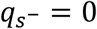 in a batch mode). 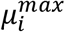 is the maximal growth rate, and 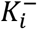 and 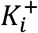 are half-saturation constants associated with the consumption of S^−^ and 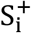.

**Kinetic parameters and stoichiometric coefficients determined through data fit:**
| Parameter | Value | Parameter | Value |
| --- | --- | --- | --- |
| $\mu_1^{\max}$ [1/h] | $2.961 \times 10^{-1}$ | $Y_{S^-/X_1}$ [g/OD] | 1.372 |
| $\mu_2^{\max}$ [1/h] | $1.658 \times 10^{-1}$ | $Y_{S^-/X_2}$ [g/OD] | 1.773 |
| $K_1^-$ [g/l] | $3.091 \times 10^{-4}$ | $Y_{S_1^+/X_1}$ [mg/OD] | 1.550 |
| $K_2^-$ [g/l] | $3.923 \times 10^{-4}$ | $Y_{S_1^+/X_2}$ [mg/OD] | 2.961 |
| $K_1^+$ [mg/l] | $3.300 \times 10^{-3}$ | $Y_{S_2^+/X_1}$ [mg/OD] | 1.365 |
| $K_2^+$ [mg/l] | $3.881 \times 10^{-4}$ | $Y_{S_2^+/X_2}$ [mg/OD] | 2.994 |
| $k_{d,1}$ [1/h] | $1.206 \times 10^{-4}$ | $q_{S^-}$ (for 3 FBs) [g/l/h] | $5.580 \times 10^{-2}$ |
| $k_{d,2}$ [1/h] | $4.024 \times 10^{-7}$ | $q_{S^-}$ (for 5 FBs) [g/l/h] | $9.300 \times 10^{-2}$ |

While the overall performance of the kinetic model was satisfactory, we found that our experimental setups were not ideal for accurately determining all model parameters. For example, **Fig. 2A** and **2B** did not effectively show the dependence of the growth of ΔTrp and ΔTry strains on tryptophan and tyrosine because the range of initial concentrations of amino acids (from 10 to 40 mg/l) was too high compared to the half-saturation constants 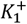 and 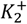 (which were determined to be 0.0033 mg/l for tryptophan and 0.00039 mg/l for tyrosine, respectively). This mismatch is partly due to the difficulty in identifying the magnitudes of half-saturation constants in advance before being determined through data fit.

### Variation in microbial interactions driven by the switch in limiting substrates in batch cultures

Based on the stochiometric and kinetic parameters determined through data fit in **Table 1**, we were able to determine microbial interactions and their variations as a function of environmental conditions using KIDI. We first analyzed various co-culture scenarios in batch modes (**Fig. 3**). In **Fig. 3A**, we considered the growth of X_1_ and X_2_ on relatively high and low initial concentrations of S^−^ (2 g/l) and 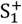 and 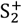 (i.e., 1 mg/l for both) as a reference condition. In the present setting, the relationship between X_1_ and X_2_ is expected to be mostly cooperative (because S^−^ is excessive in the beginning) and become competitive as the level of S^−^ decreases. While this overall trend was captured well by our model, the simulation results showed more intricate dynamics than our expectations. As depicted in the top panel of **Fig. 3A**, the concentrations of 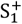 and 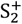 show a decreasing and increasing trend over time, respectively. This means that X_2_ does not supply sufficient 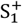 for X_1_, whereas X_1_ provides an excess of 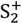 for X_2_. As a result, the growth pattern of X_1_ exhibits two distinct exponential phases. Notably, the second phase commences with a slower growth rate when 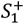 becomes depleted in the medium (the second panel from the top). This dynamic accounts for the observed slowdown in the consumption rate of S^−^ (in the first panel). Overall, these results indicate that the growth of X_1_ has a greater dependency on X_2_ than X_2_ has on X_1_, particularly when 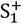 becomes depleted. This aspect is correctly captured by the higher values of *γ*_1,2_ than *γ*_2,1_ as shown in the third panel from the top. Actual values of interaction coefficients can be seen from *a*_*i,j*_, 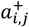, and 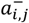 (three bottom panels of **Fig. 3A**), which also showed that 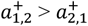 (and consequently *a*_1,2_ > *a*_2,1_) in the second growth phase of X_2_.

**Fig. 3.**
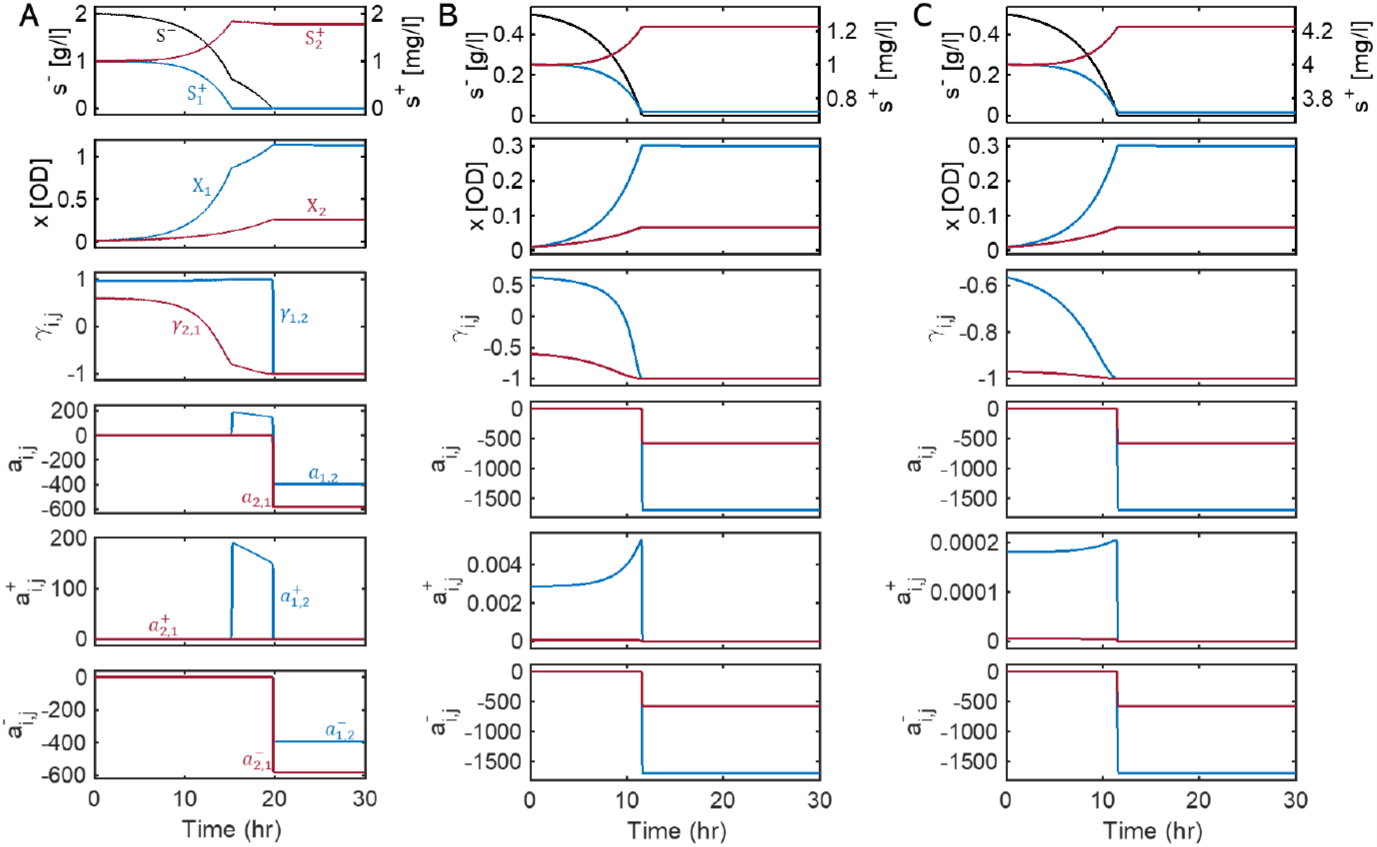
Inference of dynamic variations of interaction parameters 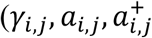 and 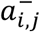 for the two *E. coli* mutants (X_1_ and X_2_) co-growing in three batch cultures. Initial substrate concentrations: **A**. 2 g/l of glucose, 1 mg/l of tryptophan, and 1 mg/l of tyrosine; **B**. 0.5 g/l of glucose, 1 mg/l tryptophan, and 1 mg/l of tyrosine; and **C**. 0.5 g/l of glucose, 4 mg/l of tryptophan, and 4 mg/l of tyrosine. Black line denotes simulated concentration of glucose (S^−^); the lines in blue and red indicate the variables and parameters associated with X_1_ and X_2_, respectively.

For comparison, we analyzed two additional conditions. (1) with a lower initial concentration of S^−^ (i.e., 0.5 g/l) (**Fig. 3B**) and (2) with lower and higher initial concentrations of S^−^ (0.5 g/l) and 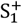 and 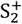 (i.e., 4 mg/l for both) (**Fig. 3C**), respectively. Unlike the first case in **Fig. 3A**, the growth profile of X_2_ does not show a biphasic growth because 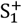 and S^−^ are depleted almost at the same time. In the case of lowering the initial concentration of S^−^ (**Fig. 3B**), KIDI showed that the level of initial competition increases (due to the limited availability of S^−^) as indicated by relatively lower values of *γ*_1,2_ and *γ*_2,1_ compared to the case of **Fig. 3A**. Notably, *γ*_2,1_ showed negative value throughout the co-growth (indicating the dominance of negative influence of X_1_ on the growth of X_2_). In the case of increasing the initial concentrations of 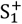 and 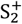 in addition to lowering S^−^ (**Fig. 3C**), the relationship between the two strains became even more negative (i.e., both *γ*_1,2_ and *γ*_2,1_ are negative), which was also an expected outcome because metabolic dependence between X_1_ and X_2_ will accordingly reduce when they can acquire what they need from the environment, rather than from partners.

In all of these cases, the relations between the two *E. coli* strains were shown asymmetric, i.e., 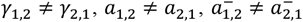, and 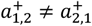 . Asymmetric interactions in terms of *a*_*i,j*_′s can also be seen over smaller time windows in **Fig. S1**. KIDI predicted *a*_1,2_ > *a*_2,1_ for the first two cases (**Fig. S1A and S1B**) but *a*_1,2_ < *a*_2,1_ for the third case (**Fig. S1C**). In the reference condition where glucose is excessive (so 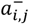’s are relatively negligible), it is mostly due to 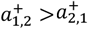 (i.e., X_1_ has a higher comparative advantage in exchanging amino acids with X_2_ than the other way around) that leads *γ*_1,2_ > *γ*_2,1_ (as well as *a*_1,2_ > *a*_2,1_) (**Fig. 3A** and **Fig. S1A**). A similar trend (i.e., *γ*_1,2_ > *γ*_2,1_ and *a*_1,2_ > *a*_2,1_) is observed in the second case where all substrates (glucose and amino acids) are limitedly available in the enviroment and therefore both 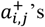 and 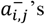 make comparable contributions to the net interaction coefficients (i.e., *a*_*i,j*_’s) (**Fig. 3B** and **Fig. S1B**). In contrast with the first two cases, the net interaction coefficients are shown to be *a*_1,2_ < *a*_2,1_ for the third case where the glucose level is low while amino acids are abundant (so 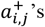 are negligible) because the magnitudes of 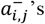 are greater than 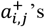. Interestingly, KIDI predicted *γ*_1,2_ > *γ*_2,1_ (**Fig. 3C**) despite *a*_1,2_ < *a*_2,1_ (**Fig. S1C**), which can happen because the implications of *γ*_*i,j*_ and *a*_*i,j*_ are not necessarily identical. The former denotes the relative dominance between promotion vs. inhibition in the relationship of species *i* with species *j*, while the latter represents the net effect of species *j* on the growth of species *i*.

### Dynamic response of microbial interactions to environmental perturbations during growth

We extend our analysis to semi-batch cultures that are perturbed by the addition of glucose FBs during growth and therefore are expected to show more dramatic changes in interspecies interactions and community dynamics. In contrast with the batch cultures considered in the previous section where no further growth is possible after the depletion of the initially added S^−^, the two strains continue to grow in semi-batch cultures due to slow but continual provision of S^−^ from the added FBs. Despite a general expectation that the competition level between the two strains will be mitigated at least at the moment of FB addition, it is uncertain: (1) to what degree this will occur under different environmental conditions, and (2) how governing microbial interactions will shift (between competition and cooperation), particularly in a later phase when the growth of the two strains is be limited by both S^−^ and 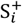. To answer these questions, we applied KIDI to the following three cases. For simplicity, we set the initial conditions to be the same as before.

First, we considered the initial concentrations of 2 g/l for S^−^ and 1 mg/l for 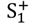 and 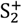 and added 3 FBs of S^−^ at around 7.5 hours (**Fig. 4A**). Due to the relatively high concentration of S^−^, the impact of adding 3 FBs of S^−^ on interactions was minimal. The profiles of interaction parameters 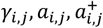and 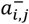 in the four bottom panels in **Fig. 3A**) as well as the growth curves of X_1_ and X_2_ showed no qualitative differences from the batch case (**Fig. 3A**), while the concentration profile of S^−^ showed an appreciable increase at the time of addition of 3 FBs (the top panel in **Fig. 3A**).

**Fig. 4.**
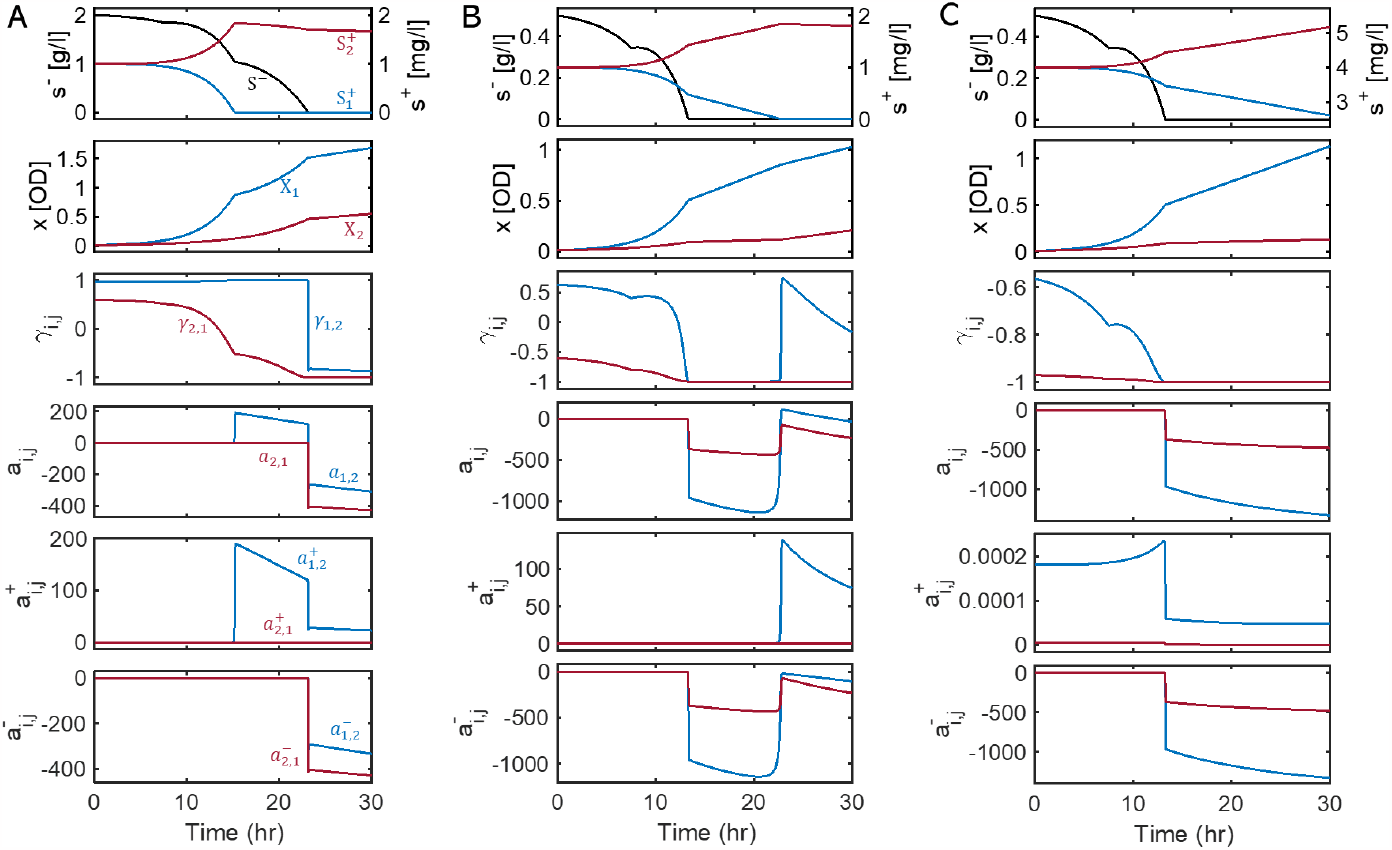
Inference of dynamic variations of interaction parameters 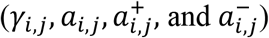 for the two *E. coli* mutants (X_1_ and X_2_) co-growing in three semi-batch cultures with 3 glucose FBs added at 7.5 hours. Initial substrate concentrations: **A**. 2 g/l of glucose, 1 mg/l of tryptophan, and 1 mg/l of tyrosine; **B**. 0.5 g/l of glucose, 1 mg/l tryptophan, and 1 mg/l of tyrosine; and **C**. 0.5 g/l of glucose, 4 mg/l of tryptophan, and 4 mg/l of tyrosine. Black line denotes simulated concentration of glucose (S^−^); the lines in blue and red indicate the variables and parameters associated with X_1_ and X_2_, respectively.

By contrast, when the initial concentration of S^−^ was low (i.e., 0.5 g/l) (**Fig. 4B**), KIDI identified the greater impact of adding FBs on both glucose concentration and microbial interactions, as indicated by sudden increases in S^−^, *γ*_1,2_ and *γ*_2,1_. Interestingly, the value of *γ*_1,2_ shifted to negative (from positive) when the medium was depleted of S^−^, but reverted to positive upon the depletion of 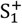. The latter suggest a substantial rise in the dependence of X_1_ on X_2_ in the absence of 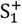 from the medium. A similar pattern was also noted for *a*_1,2_, 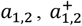, and 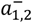 . Additionally increasing the initial concentrations of 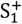 and 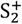 (as shown in **Fig. 4C**), thereby intensifying the level of competition, resulted in overall patterns similar to the previous case. However, both *γ*_1,2_ and *γ*_2,1_ consistently showed negative values, attributed to the heightened competition. Unlike the previous scenario, there was no increase in interaction parameters, as S_1_^+^ remained available in the medium throughout the time window up to 30 hours. Asymmetry in interaction parameters over a shorter time frame is observable in the detailed views provided in **Fig. S2**.

The simulations presented in this section demonstrate that the interactions between the two strains are highly nonlinear, influenced by the availability of S^−^, 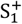 and 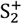 in the environment, as well as the growth characteristics of X_1_ and X_2_. As a general trend, interspecies interactions were dominated by competition when S^−^ levels were low, but shifted towards cooperation in the presence of additional limitations of 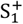 (potentially 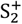 as well). Similar patterns were observed in other scenarios involving perturbations, where both the number of added FBs and the timing of their addition varied, as shown in **Fig. S3**.

## DISCUSSION

In this study, we proposed a novel computational method (KIDI) that enables quantitatively identifying environment-dependent interspecies interactions in microbial communities. By integrating growth kinetics into a gLV model, we derived an analytical form of interaction coefficients as a function of environmental variables (i.e., concentrations of chemical substrates that affect interactions), the results of which were subsequently validated through a coordinated design of co-culture experiments.

Our theoretical development significantly extends the current scope of microbial ecological modeling by completely relaxing the typical assumption of constant interactions among species. The gLV model, for example, has been widely used as a basic ecological modeling template for simulation of population dynamics and inference of interspecies interactions in microbial communities (26-28). Due to the constant interaction assumption, however, the application of the gLV model is often confined to a narrow range of conditions where interspecies interactions are expected to remain largely constant. KIDI addresses this limitation by representing interaction coefficients as an explicit function of limiting substrates. As an exception, a previous study by Momeni et al. (32) showed that pairwise interaction (i.e., gLV) models are derivable from mechanistic (i.e., kinetic) models through empirical manipulation of equations, which is however limited to special forms of kinetics and therefore cannot be generalizable (32). By contrast, our chain rule-based formulation allows us to handle any complex forms of kinetic equations with no such constraints. Consequently, KIDI enables incorporation any forms of kinetic equations as demonstrated using a double Michaelis-Menten kinetics as a demonstration example.

Dynamic variations in microbial interactions inferred by KIDI were experimentally validated using a synthetic binary consortium of two metabolically engineered auxotrophic *E. coli* mutants that cross-feed amino acids they cannot synthesize (i.e., tryptophan and tyrosine). A coordinated design of experiments provided multiple sets of data required for determining kinetic and stoichiometric parameters in the mechanistic model along with substrate concentrations, which are key inputs for quantifying environment-dependent interactions. Despite diverse culture conditions including axenic and binary growth in batch and semi-batch modes, our model with a *single* set of parameters showed satisfactory fit to the three training datasets and provided consistency with the validation dataset set aside in advance. Such a fair goodness of fit indicates the acceptability of model parameters and therefore the subsequent inference of microbial interactions. The model

Our kinetic model also shows consistency with the analysis of energetic cost of synthesizing amino acids in the literature. Mee et al. (30) estimated the energetic cost for the synthesis of 14 individual amino acids based the amounts of extracellularly supplemented amino acid and the observed growth yield of *E. coli* auxotrophic mutants. From the linear relationships between these two variables, they calculated the supplemented amounts of amino acids *per cell*, which were 1.5 ×10^7^ and 3.7 ×10^7^ for the tryptophan and tyrosine auxotrophic *E. coli* mutants, respectively. These two quantities correspond to the stoichiometric coefficients 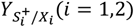 in our kinetic model, which were determined to be 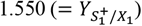 and 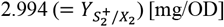 through data fit (**Table 2**). As the direct one-to-one matching between them might not be feasible, e.g., due to different units of biomass (i.e., cell number in Mee et al. (30) vs. OD in this work), we compared the ratios, which showed consistency between the two studies, i.e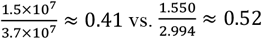.. Both results imply that compared to tyrosine, the synthesis of tryptophan is more costly. In support of this, Mee et al. (30) estimated the biosynthetic cost for tryptophan is about 43% higher than that for tyrosine.

We highlight that inferring environment-dependent interactions and their dynamic variations is a critical capability uniquely associated with KIDI. Even in a simple binary consortium considered in this work, KIDI provides new insights into interspecies interactions such as asymmetry between the two amino acid auxotrophs, which might not be obtainable otherwise. In perturbed growth experiments with glucose FBs (as in **Fig. 4B**), for example, KIDI identified that 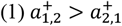 while the shared substrate (glucose) is abundant, implying that the tryptophan auxotroph (X_1_) does not support the growth of the tyrosine auxotroph (X_2_) as much as X_2_ does for 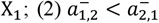 after the completion of initially added glucose until tryptophan is depleted, implying that less favorable supporters during cooperation become worse enemies when the relationship turned into competition.

While KIDI determines pairwise interaction terms following a generalized Lotka-Volterra framework, it is also capable of accounting for the influences of additional species, provided these impacts are reflected in the growth kinetics. For instance, in the case study of this article, species 1 and 2 exhibit a complex relationship, competing for glucose uptake while cooperating for amino acid exchange. However, if a third species is introduced that synthesizes and contributes amino acids to the environment more rapidly than the existing members, the dynamic between species 1 and 2 shifts. Their reliance on each other for amino acids diminishes, transforming their mixed relationship into pure competition due to the influence of the third species.

The chain rule formulation in KIDI successfully estimates interactions from given kinetics, a capability that remains effective across microbial communities of varying complexities. The primary challenge, however, is in identifying growth kinetics. This issue is especially pronounced in complex microbial communities where prior knowledge of interspecies interactions is lacking. Considering these limitations, we showcased KIDI’s effectiveness using simpler consortia, which simplify the experimental data collection needed for parameter determination in the mechanistic model. The study of model microbial consortia, extracted from natural communities, has been instrumental in enhancing our understanding of complex ecological systems (33, 34).

For KIDI to be effectively applied to simpler consortia, comprehensive measurements of all chemical and biological species involved in interspecies interactions are essential, as precise parameter identification is otherwise challenging. In our study, integrating complete temporal amino acid profiles would improve parameter identification accuracy. Typically, metabolite levels exchanged between species, such as amino acids in our case, are low and often fall below detection limits. Additional analysis of axenic culture data would help address this issue, as implemented in our work.

Despite several challenges mentioned above, it’s important to note that these stem from the difficulties in building robust kinetic models, rather than being a limitation of KIDI itself. KIDI’s primary function is to deduce the temporal variations in interaction coefficients based on environmental variables. Its unique ability to handle context-dependent interactions opens up various applications. For instance, KIDI can serve as a probing tool to investigate how assumed growth kinetics and environment-mediated mechanisms lead to specific interactions and their temporal evolution. This aspect is crucial for understanding the link between growth mechanisms of particular species and their interactions. Moreover, KIDI can greatly aid in advancing network inference techniques. The development of new algorithms for predicting microbial interactions is often hindered by a lack of benchmark data, a gap that KIDI can help fill.

KIDI’s utility goes beyond microbial ecology, encompassing a wide range of community ecology fields. This versatility comes from the fact that context dependency is not solely a microorganism trait but is also common in macro-organisms like plants and animals. For example, KIDI is applicable to classic ecological models, such as MacArthur’s consumer-resource model (35). In this model, MacArthur quantifies the impact of consumer *j* on consumer *i* (*a*_*i,j*_) based on resource population densities and associated parameters, with an underlying assumption that resource populations change more rapidly than consumer populations. KIDI is adept at deducing the competitive relationships among consumers using the chain rule, as detailed in Eq. (4) (**Supplementary Text**). This adaptability of the KIDI framework enables its extensive use in analyzing context-dependent interactions within and across different biological kingdoms in a variety of ecological systems.

## MATERIALS AND METHODS

### Mathematical definition of interaction coefficients

The dynamic change in population *i* in a community can be formulated in a general form as follows:

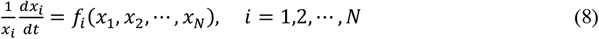

where *x*_*i*_ is the population density of species *i*, the left-hand side defines the specific growth rate of species *i*, and the function *f*_*i*_(*x*_1_, *x*_2_, ⋯, *x*_*N*_) represents a nonlinear dependence of the specific growth rate of species *i* on population densities of other species.

Using a Taylor expansion, the right-hand side of Eq. (8) can be represented as a series of polynomial terms, i.e.,

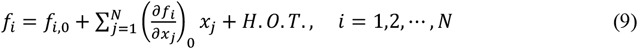

where the subscript 0 denotes a chosen reference condition, *H*.*O*.*T*. is higher-order terms. Neglecting the *H*.*O*.*T*. in Eq. (9), a gLV equation describes specific growth of species *i* using a linear equation, i.e.,

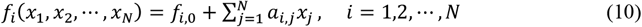

where interaction coefficient *a*_*i,j*_ denotes the effect of species population *j* on the specific growth of species *i*. For a binary community, Eq. (10) reduces to

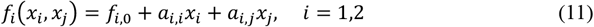

where *f*_*i*,0_ is the basal growth rate of species *i, a*_*i,i*_ is the intra-specific interaction coefficient, and *a*_*i,j*_ is the inter-specific interaction coefficient.

From Eq. (11), binary interaction coefficients in gLV are defined as follows:

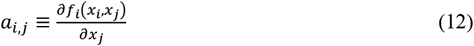

The typical formulation assumes that *a*_*i,j*_ is constant, which however leads the gLV model to fail to capture delicate dynamics of microbial interactions. Indeed, *a*_*i,j*_ is a dynamic parameter (i.e., *a*_*i,j*_(*t*)) that changes its value in varying environmental conditions as shown in the next section.

### Formulation of interaction coefficients as function of environmental variables

For simplicity, we assume in this section that *f*_*i*_ in the previous section is represented by kinetic growth rate, μ_*i*_, which is formulated as a function of nutrient concentrations in the environment. In the circumstance considered in **Fig. 1**,

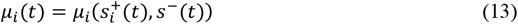

where 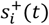 is the concentration of the nutrient (such as tryptophan or tyrosine) at time *t* that species *i* needs to get either from its partner or the environment, and *s*^−^(*t*) represents the concentration of the shared nutrient (i.e., glucose) at time *t* that two species compete for.

Based on the chain rule, we formulate *a*_*i,j*_(*t*) as a function of nutrient concentrations by plugging Eq. (13) into (12), i.e.,

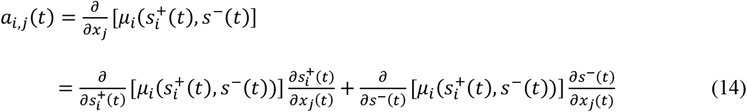

Note that the two terms on the R.H.S. of Eq. **(14)** represent the positive and negative effects of species *j* on *i* through environmental variables, i.e., 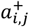 and 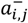 as defined below

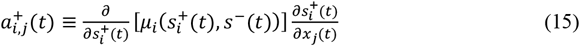

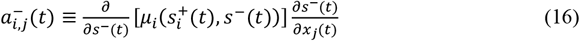

In a similar fashion, we can formulate intra-specific interaction coefficients as functions of environmental variables, i.e.,

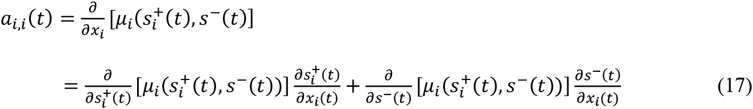

Final forms of 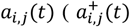 and 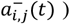 and *ai,i*(*t*) depend on specific kinetics for 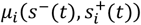. While the symbol (*t*) is dropped for simplicity, all *a*_*i,j*_’s in the main text are dynamic interaction coefficients, the values of which are changing in time as formulated in this section.

### Microorganisms and culture conditions

Two auxotrophic *Escherichia coli* (*E. coli*) mutant strains, JW2581-1 and JW1254-2 originally derived from the same strain (BW25113), were purchased from *E. coli* Genetic Stock Center (CGSC) at Yale University (http://cgsc2.biology.yale.edu/). As experimentally validated in the literature (36), these mutant strains, JW2581-1 (ΔtyrA) and JW2581-2 (ΔtrpC), are incapable of growing without supplementation of Tyrosine and Tryptophan, respectively. Each strain was incubated overnight at 37°C and 225 rpm in 50 ml of Falcon tube containing 5 ml of Lysogeny broth (LB) supplemented with 33 μg/l kanamycin. Culture cells were collected and centrifuged them at 16,000 g, 4°C for 1.5 min. The cell pellets were washed with K3 basal medium to remove residual amino acids in the samples. The washed cells were resuspended and transferred to 150 ml flasks carrying 25 ml of K3 defined minimal medium (5) containing glucose and 33 μg/l kanamycin, and cultivated at 37°C and 225 rpm. An initial absorbance at 600 nm (OD 600) was 0.04 with equivalent cell ratio. For batch mode, 4.5 g/l glucose was supplied in the culture medium. For the fed-batch mode, 0.5 g/l of an initial glucose concentration was used to shorten the lag phase and 3 or 5 glucose FeedBeads (Kühner, Basel, Switzerland), releasing glucose at constant rate, were added when OD 600 reached 0.2. We collected 500 ul of culture medium from each flask and centrifuged them at 16,000 g for 1.5 min. The supernatant and pellets were stored at -20°C until further analysis.

### Analysis of glucose concentration in the culture medium

The concentration of glucose was analyzed by a high-performance liquid chromatography (HPLC) system (Agilent, Santa Ciara, CA) equipped with a 1260 refractive index detector (RID) and an Aminex HPX-87H column (Bio-Rad, Hercules, CA). Five microliter of filtered supernatants were injected. Analytes were separated isocratically using 5 mM sulfuric acid at a flow rate of 0.7 ml/min.

### Analysis of amino acids concentration in the culture medium

The amino acids in 10 μl of filtered supernatants were analyzed using an ultra-performance liquid chromatography (UPLC) (Waters, Milford, MA) coupled with a micrOTOF II mass spectrometry (TOF-MS) system (Bruker, Bremen, Germany). Analytes were measured using a tunable UV detector at 210 and 397 nm. The amino acids were separated by an Agilent Poroshell 120 EC-C18 column at 30°C. The 1 % (v/v) of formic acid in water (mobile phase A) and 1 % (v/v) of formic acid in acetonitrile (mobile phase B) were used, respectively. The amino acids separation was obtained at a flow rate of 0.3 ml/min with a gradient program that allowed 100% of mobile phase A until 2.1 min followed by increasing mobile phase B to 40% for 2 min and then equilibrated at 0% of eluent B in a total analysis time of 6 min. Analysis of the amino acids was performed using electrospray ionization (ESI) and full-scan TOF-MS spectra (50 - 650 m/z) with 500 V end plate voltage and 4.5 kV capillary voltage. Nebulizer gas and drying gas were supplied in 1.8 bar and 8 ml/min, respectively. The dry temperature was kept at 220°C.

### Quantification of cell ratio in microbial consortium

Quantitative PCR (qPCR) was carried out in a 96-well plate by using a CFX96 Real-Time Detection System (Bio-Rad, Hercules, CA, USA). The pellets were resuspended in ultra-pure water to make consistent concentration (OD600 = 0.4) and then, the 200 μl solution was transferred to a 250 μl PCR tube. The solutions were incubated at 98°C for 10 min for cell disruption using a T100 Thermal Cycler (Bio-Rad). The lysed cells were transferred to 1.5 ml of tubes and centrifuged at 20, 000 x g for 2 min. The supernatants were analyzed by qPCR. The qPCR mixture was composed as follows: 3 μl of 10X Xtensa® buffer, 0.3 μl of primer mix (50 uM for each), 0.15 μl of i-Taq (i-DNA Biotechnology, Singapore), 3 μl of 25 mM MgCl_2_, 5 μl of purified cell lysate, and 18.55 μl of ultra-pure water. The thermal cycling was programmed as follows: 95°C for 1 min and 30 cycles of (95°C for 20 sec, 55°C for 20 sec, 68°C for 40 sec). The primers for qPCR analysis to quantify the different *E. coli* strains were provided in **Table S3**. The qPCR analysis was performed in triplicate for each sample.

### Data availability

The data and code for this study are available at https://github.com/hyunseobsong/kidi.

## ACKNOWLEDGMENTS

This work was supported by the National Research Foundation of Korea (NRF) grant funded by the Korea government (MSIT) (No. 2020R1A2C2007192), and the Korea Institute of Planning and Evaluation for Technology in Food, Agriculture, Forestry and Fisheries (iPET) funded by the MAFRA (32136-05-1-HD050). This material is also based on work supported by the National Science Foundation under Grant No. 2125155 and Nebraska Tobacco Settlement Biomedical Research Enhancement Funds to H-SS.

## SUPPLEMENTARY TEXT

### Connection between KIDI and MacArthur’s Consumer-Resource Model

In this supplementary text, we illustrate how KIDI can be extended to traditional ecological models, such as MacArthur’s consumer-resource (C-R) model, by elucidating their interconnection. MacArthur (1970) formulated the per-capita growth rate of consumer species *i* (*X*_*i*_) and resource species *j* (*R*_*j*_) as follows:

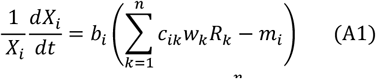

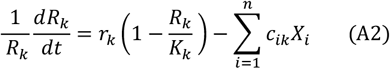

where

- *X*_*i*_ is a population density of consumer specie *i*
- *R*_*k*_ is the population density of resource species *k*
- *w*_*k*_ is the value of one unit of resource species *j* to the consumer
- *c*_*ik*_ is the rate at which consumer species *i* captures resource *j* per unit abundance of resource species *j*
- *m*_*i*_ is, the maintenance term (i.e., the total value of resource that must be harvested per capita for maintaining the current population density of consumer species *i*
- *b*_*i*_ is a factor converting the resource excess into the per-capita growth rate
- *r*_*k*_ is per-capita growth rate of resource species *j*
- *K*_*k*_ is carrying capacity for resource species *j*

MacArthur assumed faster dynamics of resource populations compared to consumer populations (i.e., *r*_*k*_ and *c*_*ik*_ are much larger than *b*_*i*_*c*_*ik*_*w*_*k*_ and *b*_*i*_*m*_*i*_) so that Eq. (A2) is reduced to an algebraic equation (by setting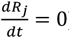). By substituting *R*_*k*_ in Eq. (A1) for *X* ′s, MacArthur derived *a*′_i*j*_ (i.e., the competitive relationship between the two consumers *i* and *j*, or more precisely the impact of consumer *j* on the growth on consumer *i*) in the Lotka-Volterra (LV) model as follows:

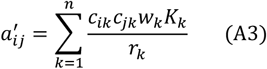

where we use the prime notation in *a*′ to differentiate it from the definition of *a*_*ij*_ commonly used in a generalized LV model. These two notations are related by 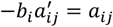.

We now show that the above equation can be derived by KIDI. Interspecies interaction coefficient *a*_*ij*_ (not 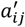) for the system with multiple resources in KIDI are defined as follows:

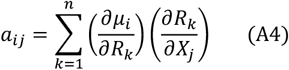

where μ_*i*_ is the specific (or per-capita) growth rate of species *i* 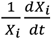. The first term 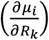 is obtained simply by taking the derivative of the per-capita growth rate for *X*_*i*_ with respect to *R*_*k*_ in Eq. (A1), i.e.,

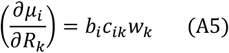

Next, we take the derivative of the steady-state equation of Eq. (A2) with respect to *X*_*j*_ to give the second term 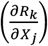 as follows:

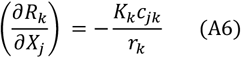

With the substitutions using Eqs. (A5) and (A6), the final form of *a*_*ij*_ becomes

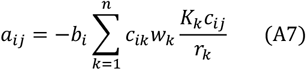

The above equation proves 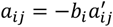.

## SUPPLEMENTARY TABLES AND FIGURES

**Table S1.** Initial culture conditions for experimental data displayed in Fig. 2.

| Exp # | Corresponding figures | Initial inoculum size (OD) | Glucose (mg/l) | Tryptophan (mg/l) | Tyrosine (mg/l) | # of FBs added | Time FBs added (hr) |
| --- | --- | --- | --- | --- | --- | --- | --- |
| 1 | Fig. 2A (top panel) | ΔTrp only: 0.01 | 5.838 | 10 | - | - | - |
| 2 | Fig. 2A (middle panel) | ΔTrp only: 0.01 | 5.870 | 20 | - | - | - |
| 3 | Fig. 2A (bottom panel) | ΔTrp only: 0.01 | 5.936 | 40 | - | - | - |
| 4 | Fig. 2B (top panel) | ΔTyr only: 0.01 | 5.920 | - | 10 | - | - |
| 5 | Fig. 2B (middle panel) | ΔTyr only: 0.01 | 5.909 | - | 20 | - | - |
| 6 | Fig. 2B (bottom panel) | ΔTyr only: 0.01 | 5.863 | - | 40 | - | - |
| 7 | Fig. 2C (top panel) | ΔTrp: 0.01<br>ΔTyr: 0.01 | 3.851 | 0.4 | 0.4 | - | - |
| 8 | Fig. 2C (middle panel) | ΔTrp: 0.01<br>ΔTyr: 0.01 | 0.626 | 0.4 | 0.4 | 3 | 7.5 |
| 9 | Fig. 2C (bottom panel) | ΔTrp: 0.01<br>ΔTyr: 0.01 | 0.620 | 0.4 | 0.4 | 5 | 7.5 |
| 10 | Fig. 2D (top panel) | ΔTrp: 0.01<br>ΔTyr: 0.01 | 2.878 | 0.4 | 0.4 | - | - |
| 11 | Fig. 2D (middle panel) | ΔTrp: 0.01<br>ΔTyr: 0.01 | 0.375 | 0.4 | 0.4 | 3 | 10 |
| 12 | Fig. 2D (bottom panel) | ΔTrp: 0.01<br>ΔTyr: 0.01 | 0.377 | 200 | 200 | 3 | 10 |

**Table S2.**
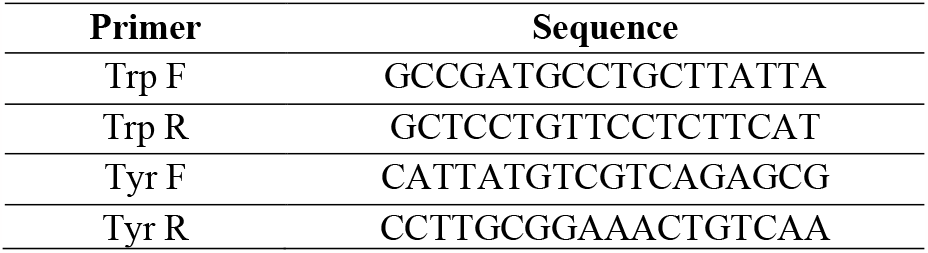
Sequence of the primers used in the qPCR analysis.

**Figure S1.**
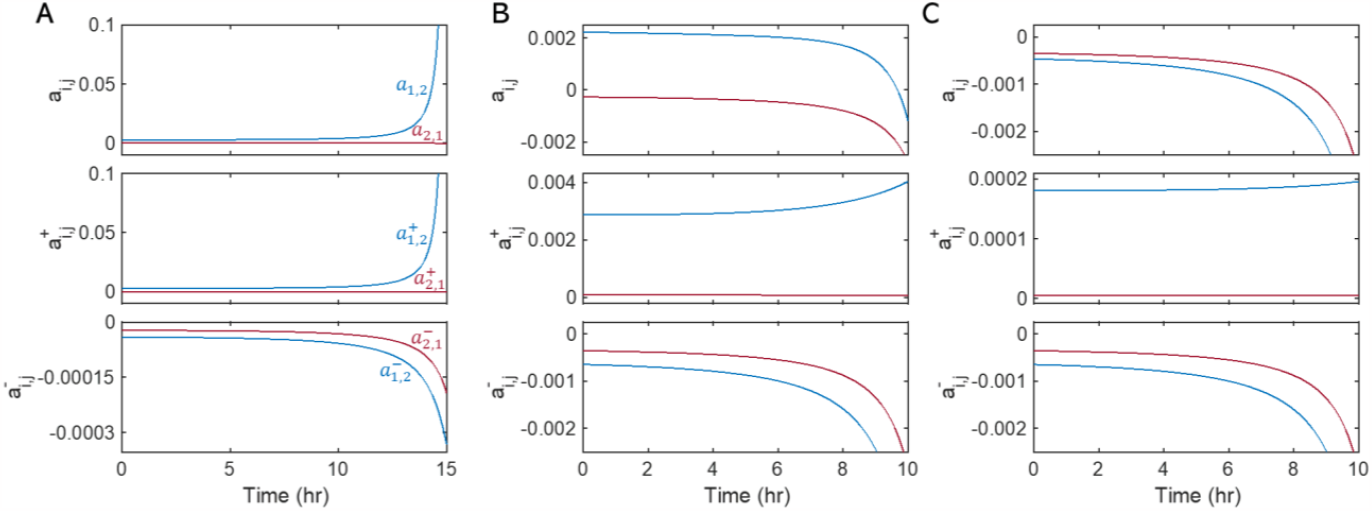
Zoom-in views of in interaction parameters, *a*_*i,j*_, 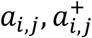, and 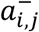 in **Figs. 3A, 3B, and 3C**, respectively. The color scheme is the same as in **Fig. 3**.

**Figure S2.**
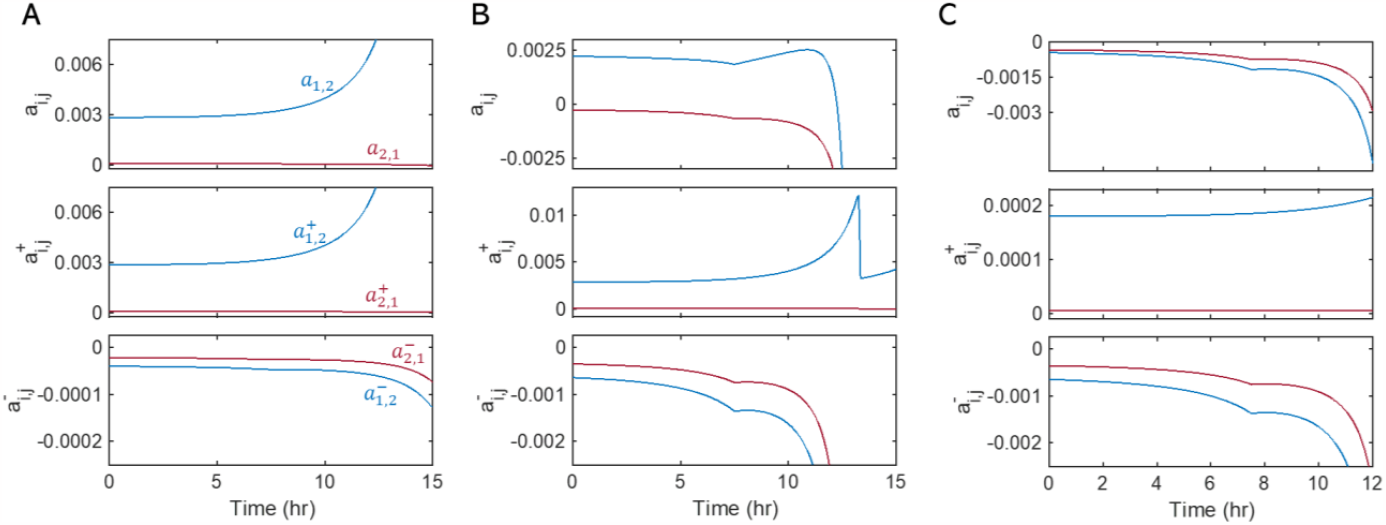
Zoom-in views of in interaction parameters, *a*_*i,j*_, 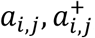, and 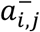 in **Figs. 4A, 4B, and 4C**, respectively. The color scheme is the same as in **Fig. 4**.

**Figure S3.**
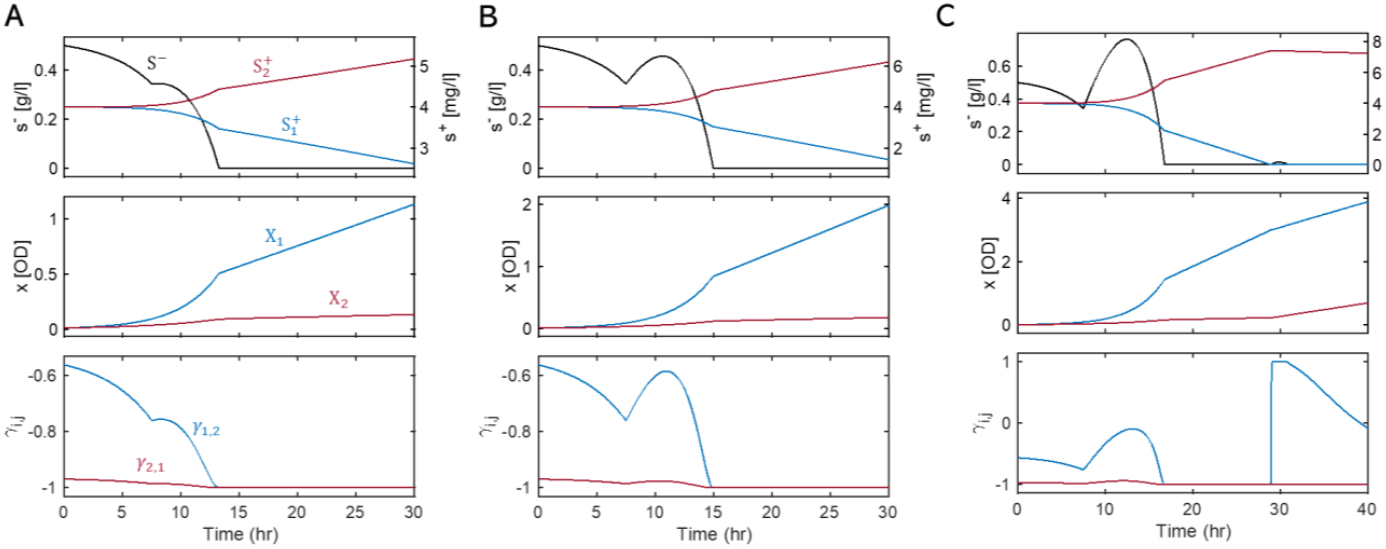
The predicted impacts of the number of added FBs on the substrate profile (S^−^), the population dynamics (X_1_ and X_2_), and the interaction parameter (*γ*_*i,j*_) in fed-batch cultures with initial substrate concentrations of 2 g/l of glucose, 0.4 mg/l of tryptophan, and 0.4 mg/l of tyrosine. The number of glucose BFs added at 7.5 hours: **A**. 3 FBs; **B**. 6 FBs; and **C**. 10 FBs.

**Figure S4.**
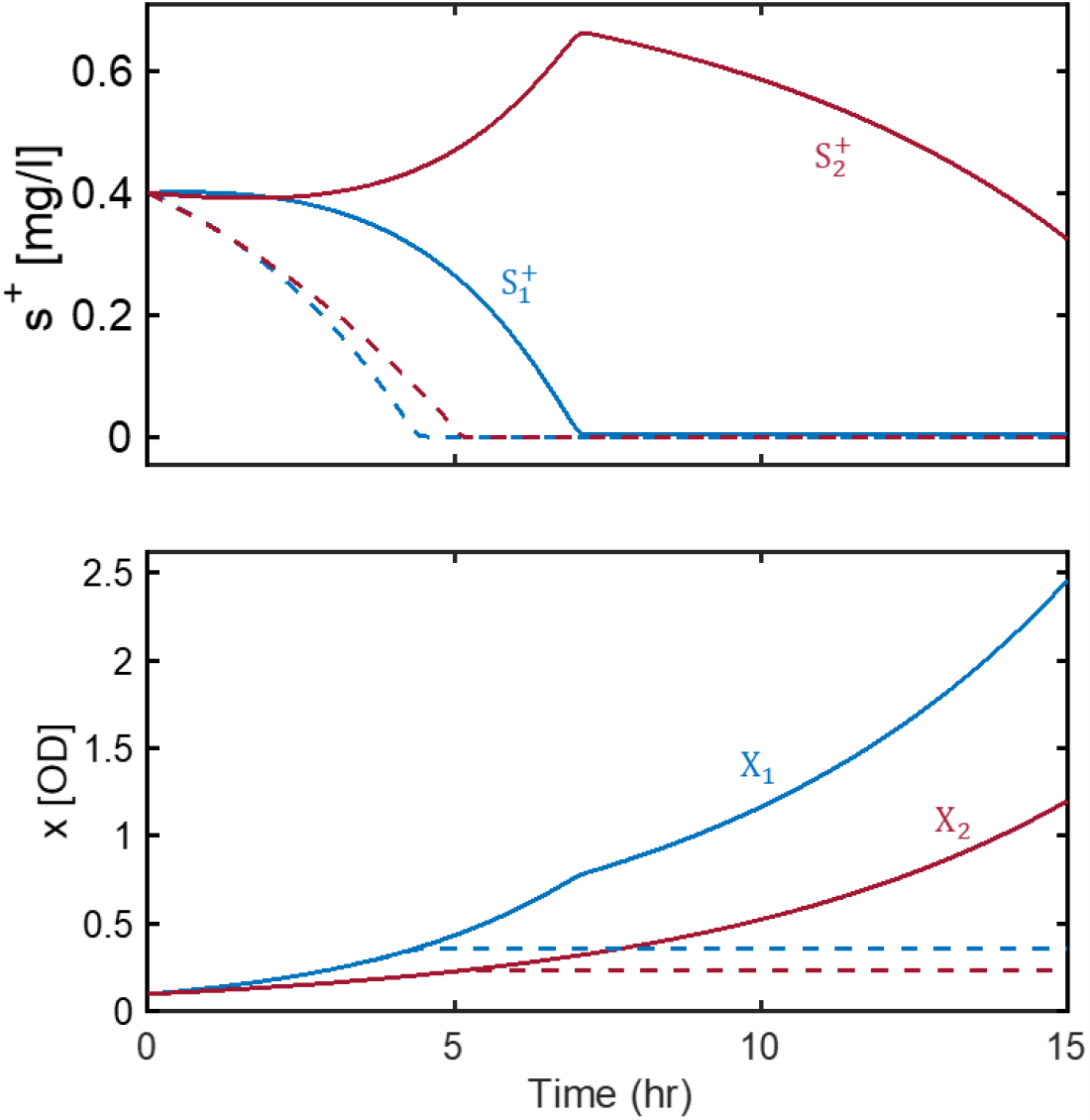
Simulated substrate concentrations (top), and biomass concentrations (bottom) with and without amino acid cross-feeding in the batch culture with excessive provision of glucose. Initial substrate concentrations of amino acids: 0.4 mg/l of tryptophan and 0.4 mg/l of tyrosine. Solid lines denote the case with cross-feeding; dashed lines denote the case of preventing cross-feeding in simulations.

